# Influence of ULK1 activity on the memory of ATG101 auto-activation dynamics

**DOI:** 10.1101/2025.06.27.661946

**Authors:** Anoshi Patel, Sonja Schneider, Gabor Nagy, Olexandr Dybkov, Henning Urlaub, Helmut Grubmüller, Alex C. Faesen

## Abstract

The initiation of autophagy is marked by the prompt convergence of initiation proteins to form super molecular complexes in spatially-defined autophagic hubs. The recruitment and activation of the ULK-kinase complex is a pivotal event in starting the initiation cascade. ULK complex-component ATG101 is required for initiating autophagy in mammalian cells, but its function is unknown. Its HORMA domain interacts with the HORMA domain of ATG13 and transmembrane protein ATG9A to collectively form the essential autophagy initiation complex ATG9A-ATG13-ATG101. ATG101 has structurally malleable elements, reminiscent of topological conversions used in related metamorphic HORMA domain proteins to control the rate of complex formation. To elucidate the function of ATG101, we monitored the interaction kinetics of ATG101 with ATG13 and ATG9A, and observed that the interaction of ATG101 with ATG9A and ATG13 is exceptionally slow. Dramatic acceleration of complex formation is observed upon a change in the fold of ATG101 induced by its transient homo-dimerization, which in turn is initiated by phosphorylation by the ULK1 kinase. In an auto-catalytic mechanism, ATG101 dimers create a positive feedback to propagate activation to further ATG101 molecules in the absence of ULK1 activity. Despite the competitive nature of the interaction interfaces, homo-dimerization of ATG101 surprisingly accelerates its association to ATG13. Memory of ATG101 activation persists for many hours after dephosphorylation and continues to accelerate the assembly of the ATG9A-ATG13-ATG101 complex. Overall, this work proposes an unusual regulatory mechanism where UKL1 initiates an ATG101 auto activation cascade, whose memory creates a responsive positive feedback that dictates the assembly rate of a key complex in autophagy initiation.

## Introduction

Macro-autophagy (called autophagy hereafter) is a conserved process of regulated degradation. It eliminates damaged and unnecessary cellular components by transporting biomolecules to lysosomes, where they are degraded to be recycled. The ULK1 complex responds to upstream nutrient and energy signals and is essential to initiate autophagy by bridging and building the autophagy initiation machinery^1^. When fully assembled, the ULK complex is composed of four proteins: the ULK1 or -2 kinase, ATG13, ATG101, and focal adhesion kinase family interacting protein of 200 kDa (FIP200)^2–5^. ULK kinase activity is a key regulatory event to initiate and coordinate the autophagy pathway^6^. For example, ULK1 phosphorylates VPS15 to activate the PI3-kinase complex^7^, but also its own components like ATG101 with unknown consequences^6^.

The other subunits of the ULK complex are thought to have numerous scaffolding and bridging roles, some of which act independently and upstream of the ULK kinase and its activity. The recruitment of ATG13 and ATG101 to sites of autophagy initiation is an essential and defining early step in both starvation-induced autophagy and selective autophagy^3,8,9^. The C-terminal 300 residues of ATG13 are predicted to be intrinsically disordered and interacts with FIP200 and the ULK kinase^10–12^. The function of ATG101 is unclear, but it has been reported to increase the lifetime of its binding partner ATG13 by protecting it from proteasomal degradation and to stabilize basal phosphorylation of ULK1 and ATG13^3,5^. Hetero-dimerization of ATG13 and ATG101 via their HORMA domains (named after the Hop1p, Rev7p and MAD2 proteins) is essential for autophagy, which might in part be due to the reciprocal stabilizing effect^13–15^. Notably, both ATG13 and ATG101 are present in higher abundance than the rest of the ULK kinase complex and might have additional roles outside the ULK kinase complex^15^. For example, an ATG13-ATG101 dimer lacking an ULK kinase assembles with the early autophagy machinery, such as ATG9A, to promote phagophore growth in fed cells^8,9,16,17^. The ATG13-ATG101 dimer interacts with the “HORMA dimer–interacting region” (HDIR) in ATG9A and recruits ATG9A during p62-dependent autophagy, independent of FIP200 and ULK1^13,16,18^. Deletion of ATG13 or ATG101 abrogates their reciprocal interactions, and leads to a mis-localization of ATG9A and phenotypes similar to an ATG9A knock-out^16,19^. The ATG13-ATG101 dimer plays a critical coordinating role in recruitment of downstream factors to the autophagosome formation site by nucleating the assembly of stable multi-megadalton complexes^3,13,15,16,20–23^. However, it is currently unclear how the assembly of these complexes is regulated in space and time.

Despite its importance, there are no known ATG101 orthologs in *S. cerevisiae*, but it is present in lower metazoans. Human ATG101 (218 amino acids), does not have obvious sequence similarity to any other protein in species that lack this gene. The X-ray structures of human ATG101 show that its fold is promiscuous. ATG101 in isolation adopts a similar fold as the ‘open’ conformer of the prototypical metamorphic HORMA domain protein MAD2 as defined by the position of the C-terminus into a beta-sheet^24^ (**Figure 1A**). The fold of ATG101 does however change upon dimerization with ATG13, which includes the ejection of the C-terminus and its conversion from a beta-strand into an alpha-helix^22,24^ (**Figure 1A**). This topological conversion allows the interaction of the newly formed C-terminal helix with the PI3-kinase complex^22^. Together, the ATG13-ATG101 hetero-dimer is analogous to the ‘open’-‘closed’-conformational homo-dimer of MAD2^25^. The topologically asymmetric MAD2 homo-dimer is an obligatory reaction intermediate in the mechanism to accelerate the assembly of the Mitotic Checkpoint Complex, by lowering the activation barrier of the otherwise very slow metamorphosis of ‘open’ MAD2 to the ‘closed’ conformer^26,27^. Therefore, the default ‘open’ conformer of MAD2 represents an autoinhibited state and the dimerization accelerates the rate-limiting conversion into the active ‘closed’ conformer in order to induce the formation of its downstream effector complex. The emerging paradigm is that regulated acceleration of metamorphosis of HORMA domain proteins controls the rate of signalling or assembly of effector complexes^27,28^. This paradigm might be conserved beyond HORMA domain proteins, as dimerization has been found to modulate interaction kinetics of unrelated metamorphic proteins^25,29–32^. Using qualitative interaction assays, we had previously shown that the interaction kinetics of ATG13 and ATG101 to ATG9A are remarkably slow, similar to related HORMA domain proteins MAD2 and REV7^13,26,27,33,34^. This suggested that both ATG13 and ATG101 too default to an autoinhibited state. Indeed, ATG13 adopts two distinct folds that could be identified and separated based on their differential surface charge and mutually exclusive interactions^13^. Mutants designed to interfere with the metamorphic behaviour in ATG13 and ATG101 both show strong autophagic defects and do not allow for the assembly of the initiation machinery^13^.

**Figure 1:**
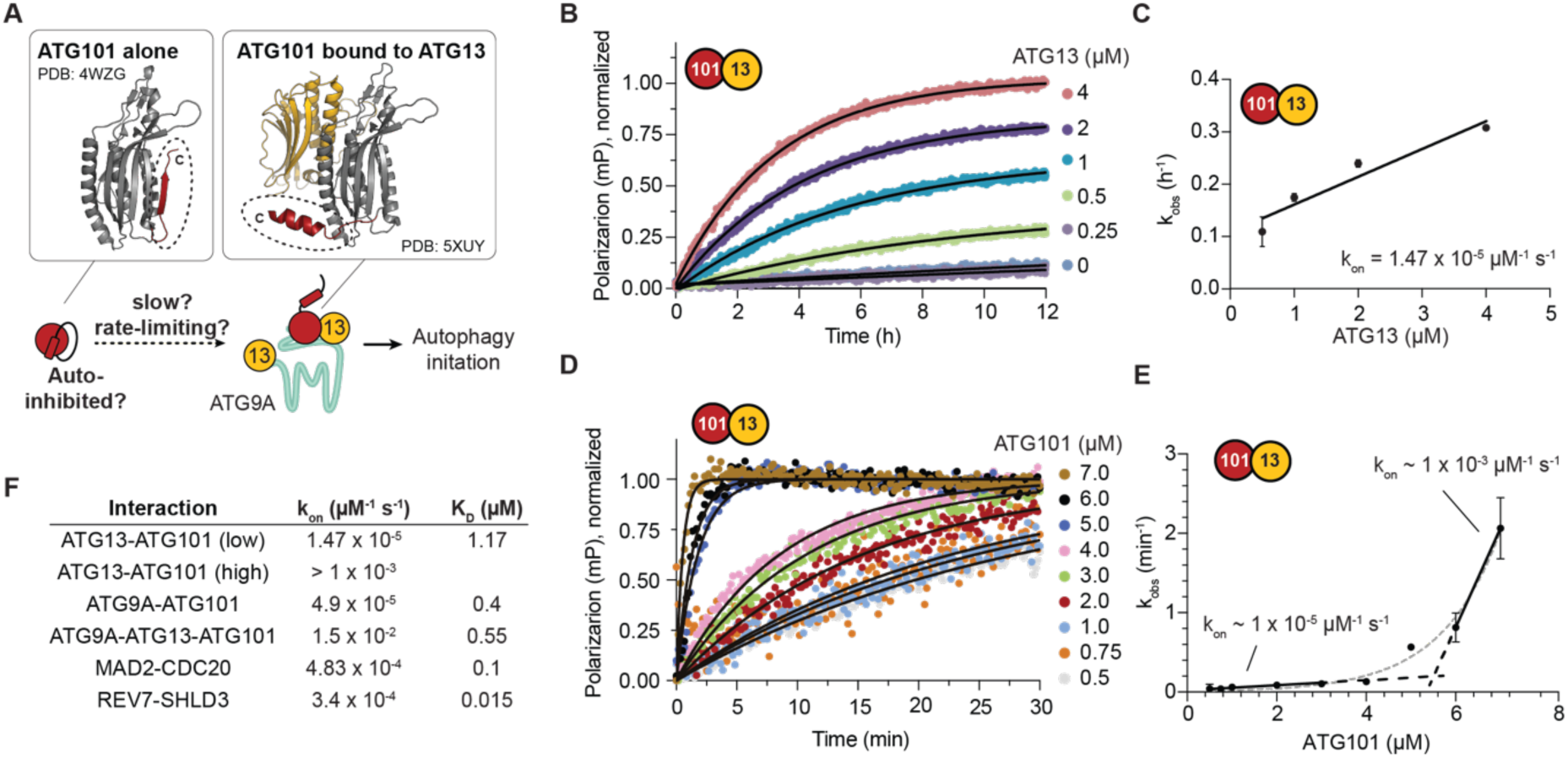
The interaction of ATG101 with ATG13 is exceedingly slow, but suddenly accelerates at higher ATG101 concentration. **a)** The structures of ATG101 suggest an activation might be required to interact with ATG13 and ATG9A in order to initiate autophagy. **b,c)** The interaction of ATG101 with ATG13 is exceedingly slow. Time zero is the first time point after mixing 20 nM ATG101^Alexa488^ with indicated ATG13 concentrations. All panels in **b** reporting time-dependent changes in Fluorescence Anisotropy signal are single measurements representative of at least three independent technical replicates of the experiment. After single exponential fitting of the curves in **b**, the apparent first order rate constants (k_obs_) were plotted as function of ATG13 concentration in **c**, with k_on_ being the slope of the resulting line. **d,e)** The slow association of ATG101 to ATG13 accelerates at least two orders of magnitude depending on ATG101 concentration. Fluorescence Anisotropy measurements as performed in **b** and **c**, but instead mixing 50 μM unlabeled ATG13 with indicated ATG101 concentrations.

The role of ATG101 is enigmatic, in particular the contribution of its topological conversion to its function and interaction kinetics. In this study, we monitored the interaction kinetics of ATG101 in real time at physiologically relevant concentrations using an *in vitro* Fluorescence Polarisation (FP) sensor. We observed that the interactions of ATG101 with either ATG13 and ATG9A have remarkably small on-rate constants, which are however dramatically accelerated at increased concentrations of ATG101. We show this is due to transient homo-dimerization of ATG101, which is stabilized after phosphorylation by ULK1, leading to the activation of ATG101 and rapid assembly of the ATG9A-ATG13-ATG101 complex. This mechanism results in a memory of activation, as ATG101 ‘remembers’ its activation for many hours after dephosphorylation. This suggests that the initial phosphorylation induces a structural conversion into an active state required for dimerization. Indeed, shifts in CD spectra indicate structural changes correlating with the topological conversion of the beta strand into an alpha helix. Although it paradoxically uses the same interaction interface to engage with ATG13, the transient nature of the homo-dimerization promotes, rather than inhibits, the interaction of ATG101 to ATG13 and subsequent interaction with ATG9A. Moreover, only a small amount of activated ATG101 suffices to auto-catalytically activate all ATG101 molecules to accelerate the formation of the key ATG9A-ATG13-ATG101 complex. Overall, this suggests an unusual regulatory mechanism, where the ATG101 dimer serves to present a template for the conversion of subsequent ATG101 molecules to create a responsive positive feedback and initiate the local fast assembly of the autophagy machinery on demand.

## Results

### The interaction of ATG101 with ATG13 and ATG9A is exceedingly slow

Structures of ATG101 show a striking difference in the topology of its C-terminal 25 residues. Alone, this region folds as a beta-strand incorporated into an extended beta sheet^24^. However when bound to ATG13, the terminal beta-strand is dislodged from the extended sheet to form an alpha-helix^22,24^ (**Figure 1A**). This C-terminal mobile structural element corresponds with the topologically mobile ‘seat-belt’ in MAD2, which is elementary in defining its metamorphic state and thus interaction spectrum^25,35^. Structure prediction software has a mixed track record in predicting multiple metamorphic conformers from single polypeptides^36–38^. For example, AlphaFold3^39^ is somewhat uncertain about the C-terminal ATG101 region, but changes its prediction from the beta-strand to the helical conformer upon introducing the ATG9A HDIR peptide or ATG13 HORMA domain (**Figure S1A**). We note that the C-terminal region is missing in the ATG13-ATG101-ATG9A^HDIR^ structure and therefore does not guide the prediction^18^. This structural conversion is allosterically induced, as this region is not part of, nor close to, the ATG13-ATG101 dimer interface.

Since the C-terminal region is not part of the interaction interfaces and seems to prefer to fold as a beta strand by default, it is therefore unclear what comes first: the hetero-dimerization with ATG13 or the conversion to the alpha-helix? In case of the latter, the prediction would be that the ATG13-ATG101 heterodimer would form slowly due to significant unfolding of ATG101 required, and thereby create a rate-limiting step in complex formation (**Figure 1A**). To test this, we developed an *in vitro* fluorescence polarization (FP) assay by fusing an Alexa488 moiety to the C-terminus of ATG101 using Sortase labeling^40^ (**Figure S1B**). This allowed us to determine binding isotherms and quantify apparent binding strengths (*K_D,app_*) of ATG101 to ATG13, ATG101 to ATG9A and ATG13-ATG101 to ATG9A, respectively (**Figure S1C-E**). This sensor also allowed us to monitor ATG13-ATG101 complex formation in real time (**Figure 1B**). The time-dependence increase of the FP signal could be fitted with a single exponential function to yield an apparent reaction rate (*k*_obs_) that increased linearly with the concentration of ATG13, indicative of pseudo-first-order kinetics (**Figure 1C**). Even at above physiological concentrations of ATG13 (500 nM, using 20 nM ATG101), complex assembly is very slow, with a half-life (*t*_1/2_) of approximately 13 hours. The corresponding association rate of 1.47 × 10^-5^ μM^-1^s^-1^ is several orders of magnitude slower than typical protein-protein interactions, and an order of magnitude slower than other known metamorphic HORMA domain proteins MAD2^27^ and REF7^33^, highlighting the extraordinary slow nature of the assembly of the ATG13-ATG101 complex (**Figure 1C,F**). The association of ATG101 with ATG9A was equally slow, indicating that likely an inhibitory, and thus rate limiting, event in ATG101 prevents a canonical interaction to both ATG13 and ATG9A (**Figure S1F,G**).

Importantly however, when the ATG13-ATG101 complex was allowed to pre-form during an overnight pre-incubation, the interaction with ATG9A was complete within 1 minute at near-physiological concentrations (125 nM) (**Figure S1J**). The pre-incubation accelerated the interaction to ATG9A by 3-orders of magnitude to 1.5 × 10^-2^ μM^-1^s^-1^ (**Figure S1K**). Overall, this suggests that the ATG13-ATG101 dimerization is the limiting step for association with ATG9A and that the interaction with ATG13 lifts the inhibitory effect in ATG101 that prevents prompt complex formation.

### Concentration controls the unusual activation of ATG101 interaction dynamics

The rate-limiting nature of the ATG101 interaction kinetics prompted us to invert the experiments, and instead vary the concentration of ATG101. We used the FP assay to observe the interaction kinetics with ATG13 or ATG9A at varying ATG101 concentrations. This yielded two surprising, and at first glance contradictory, observations. Most prominently, we observed that ATG13-ATG101 complex formation was slow until it suddenly accelerated at concentrations above ∼5 μM ATG101, taking place within a few minutes (**Figure 1D**). This was corroborated by quantifying the on-rates, where the typical linear increase of rates is followed by sudden atypical acceleration at elevated ATG101 concentrations (**Figure 1E**). This switch-like 100-fold increase of on-rate could therefore not be explained by the increase of ATG101 concentration alone. Conversely, we observed that the interaction of ATG101 with ATG9A, in the absence of ATG13, slows down with increasing ATG101 concentration (**Figure S1L,M**). These results suggest that the association of ATG101 with its binding partners follows an unusual mechanism, which has an obligatory intermediate that is sensitive to the concentration of ATG101 and directly controls both interaction dynamics and strength. The inhibited interaction of ATG101 to ATG9A shows the order of interaction: the formation of the ATG13-ATG101 dimer has to precede the interaction of ATG101 with ATG9A.

### ATG101 transiently homodimerizes

These observations indicate that there is an inhibitory element in ATG101 that prevents a straightforward interaction to ATG13, and that this inhibition is overcome at increased concentrations of ATG101. Metamorphic proteins are known to modulate their interaction kinetics by inducing the transition from a (default) autoinhibited-conformer to an interaction competent-conformer via homo-dimerization^25,29–32^. We have not observed homodimerization of ATG101 during any size exclusion chromatography experiment (using wild-type, nor modified protein (see below)), although the elution peaks were typically not symmetric, suggesting heterogeneity (**Figure S1B**). Regardless, we performed a pull-down experiment using differentially tagged ATG101 fusions at 2 or 6 μM (bait and prey, respectively) concentration, and using quick washes with reduced volumes to keep the concentration relatively elevated. This approach revealed that ATG101 can indeed stoichiometrically homo-dimerize at concentrations that accelerate ATG13-ATG101 complex formation (**Figure 1E** and **2A**). This dimer uses the canonical HORMA dimerization interface, as the introduction of the L29R and H30R mutations^41^ abrogated both the ATG101 homo-dimerization as well as the hetero-dimerization with ATG13 (**Figure 2B,C**). As we observed a decreased binding to ATG9A at elevated ATG101 concentrations, we wondered if the ATG101 homo-dimer would be incompatible with ATG9C binding. Indeed, when fusing ATG101 with a GST-tag to promote homodimerization, the interaction with ATG9A was strongly reduced compared to an MBP-tagged ATG101 (**Figure 2D**).

**Figure 2:**
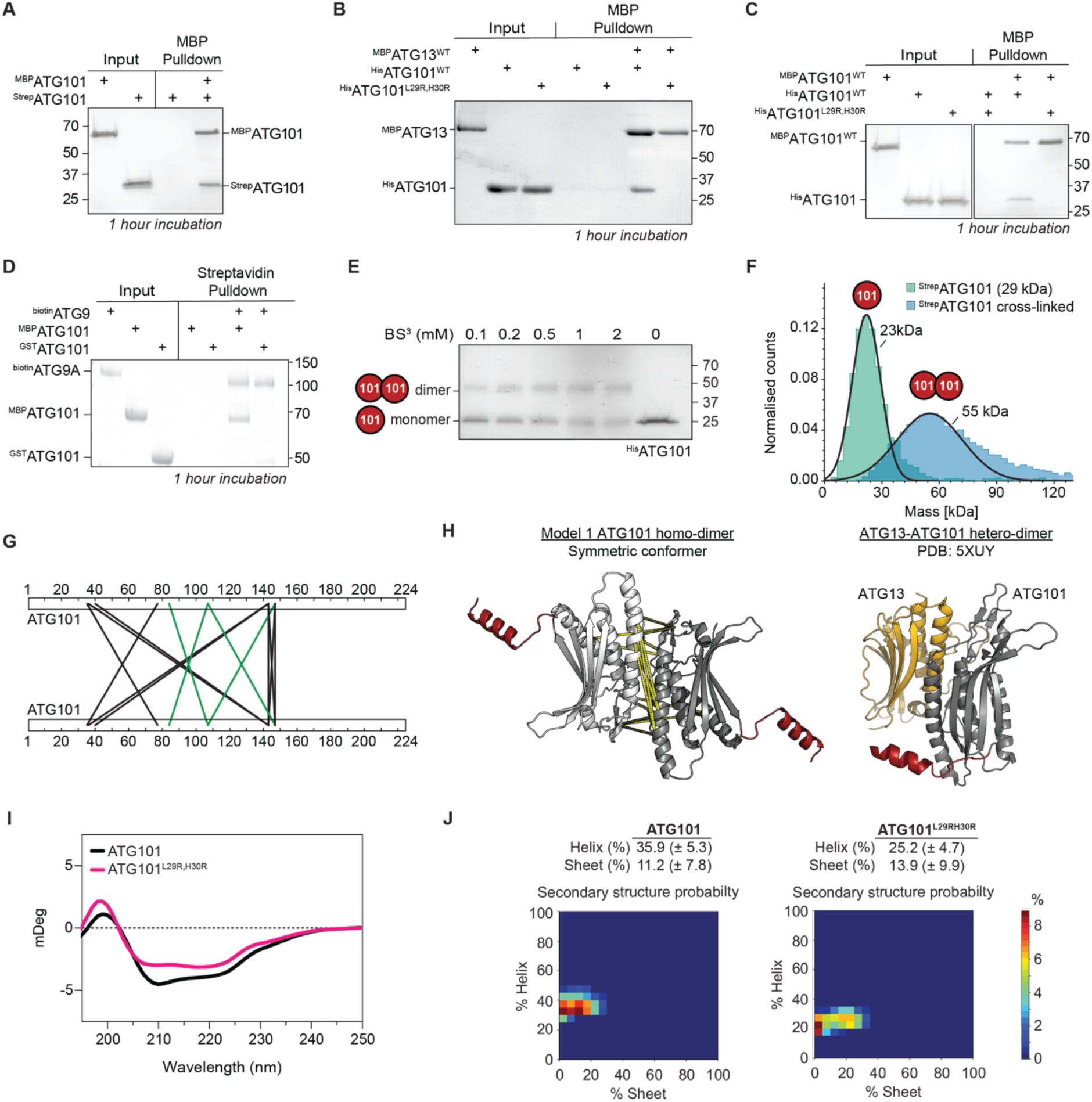
ATG101 transiently homo-dimerizes using the canonical HORMA dimer interface. **a)** ATG101 can homo-dimerize. Pull-down experiment using 2 μM MBP-ATG101 as bait and 6 μM Strep-ATG101. **b,c)** ATG101 interaction with ATG13 (panel **b**) and homo-dimerization (panel **c**) require the same interface. Pull-down experiments using 2 μM MBP-ATG13 (panel **b**) or 2 μM MBP-ATG101 (panel **c**) as bait and 6 μM His-ATG101. Introduction of the L29R and H30R (reference 41) mutations abrogated both the homo-as well as the hetero-dimerization with ATG13. **d)** Inducing homo-dimerization via a dimeric GST-fusion prevents the interaction of ATG101 to ATG9A. Pull-down experiment using 1 μM ATG9A with biotinylated amphipol as bait and 3 μM GST- or MBP-ATG101. Quantification in **Fig S2B** of three independent technical replicates of the experiment. **e)** Incubation of ATG101 with cross-linker BS^3^ results in the appearance of a distinct second protein with a mass corresponding to an ATG101 dimer. Input is shown in lanes with no BS3 (0 mM). **f)** ATG101 homo-dimerization is stabilized by crosslinking. Mass photometry experiment using ATG101 (150 nM), comparing untreated and after BS^3^ (1 mM) treatment and subsequent purification. **g,h)** Crosslinking with BS^3^ coupled with mass spectrometry confirmed that ATG101 homo-dimerizes via the canonical HORMA dimer interface. The cross-links determined in **g** were used to predict three structural models using HADDOCK^42^ in **h** and **S2C** of the ATG101 homo-dimer. The resulting models show that ATG101 uses the same interface for both homo- (**h,** left) and hetero-dimerization (**h,** right). Cross-linked residues in the dimer interface are shown as black in **g** and yellow beads on string in **h**). **i,j)** Dimerization of ATG101 induces structural changes. CD spectra (panel **i**) recorded using 3.5 μM ATG101 or the ATG101^L29R,H30R^ dimerization defective mutant, where analysed and modelled using SESCA^44,45^ (panel **j**), where the proposed models show an increase in helical content and concomitant decrease of beta-sheets in ATG101 upon dimerization.

Since ATG101 homo-dimerization, in contrast to ATG13-ATG101 hetero-dimerization, seems to be weak and/or transient, we aimed to stabilize it using the crosslinker BS^3^. In the presence of the crosslinker, we observed the emergence of a single additional protein species with a mass corresponding to that of a dimer when using the wild-type protein, but not when using the ATG101^L29R,H30R^ dimerization-deficient mutant (**Figure 2E** and **S2A**). Using mass photometry, we confirmed that this treatment stabilized the homo-dimer (**Figure 2F**). We determined the sites of cross-linking using mass-spectrometry (**Figure 2G** and **Table S1**) and used these to create a model of the dimer using HADDOCK^42^. The cross-links agreed well with a model that ATG101 homo-dimerization uses the canonical HORMA dimer interface (**Figure 2H** (left) and **S2C**). The model thus highlighted that homo-dimerization is competitive with the interaction with ATG13 (**Figure 2H**, right).

The cross-links cannot distinguish if the dimerization is symmetric (meaning with specifically the same conformer, as seen with REV7 in PolZ^43^), asymmetric (specific for opposing conformers, as seen for MAD2^25^), or agnostic (there is no selectivity, as seen for REV7 in Shieldin^33^). Regardless, as dimerization typically stabilizes the change in fold in metamorphic proteins, we wondered if the same is true in ATG101. Therefore, we used circular dichroism (CD) measurements of ATG101 and the ATG101^L29R,H30R^ mutant, which showed significant changes in secondary structure upon dimerization (**Figure 2I**). We used SESCA^44,45^ which uses a Bayesian statistics approach to determine the likelihood of possible secondary structures of a target protein based on the measured CD spectra (**Figure 2J**). The proposed models show an increase in helical content and concomitant decrease of relative beta-sheets in ATG101 upon dimerization. These changes agree with the observed conversion of the C-terminal beta-strand into an alpha-helix in the published ATG101 structures (**Figure 1A**). Overall, this shows that dimerization initiates a structural and topological conversion in ATG101.

### ULK1 activity induces ATG101 structural conversion and homodimerization

Homo-dimers of metamorphic proteins usually contain at least one converted or intermediate ‘unbuckled’ conformer^25,29–32^, therefore we wondered if structural changes in ATG101 are required to induce homo-dimerization. Residues Ser11 and Ser203 of ATG101 are known phosphorylation sites of ULK1 with unknown function^6^. Ser203 is part of the C-terminal topologically flexible region, and could hypothetically directly destabilize the beta-strand and thereby promote the formation of the converted conformer (**Figure 1A**). Indeed, AlphaFold3 changes its prediction to the helical conformer upon introducing the phosphorylation at Ser203 (**Figure S1A**). To test if phosphorylation affects ATG101 structure and its ability to homo-dimerize, we added the generic phosphatase inhibitor Okadaic Acid during the last 5 hours (of the insect cell culture expressing ATG101, resulting in the purification of phosphorylated protein (**Figure S3A**). Using mass-photometry we confirmed that this phosphorylation results in ATG101 dimerization in contrast to unphosphorylated ATG101 at 150 nM (**Figure 2F** and **S3B**). Next, we co-expressed ATG101 and ULK1 by mixing their respective baculo-viruses (in a 5:1 ratio) to the expression culture, which also resulted in the phosphorylation and dimerization of purified ATG101 (**Figure 3A** and **S3C**). This is a direct result of ULK1 activity, as the phosphorylation of ATG101 using purified full-length recombinant ULK1 kinase yielded an ATG101 homo-dimer with an apparent Kd between 100 and 250 nM (**Figure 3B,C**). Using mass-spectrometry, we confirmed that ATG101 is indeed phosphorylated by ULK1 at Ser11 and Ser203 (**Table S2**). A similar stabilization of the ATG101 homo-dimer was observed using the Ser11 and Ser203 phospho-mimetic mutants (**Figure S3D**).

**Figure 3:**
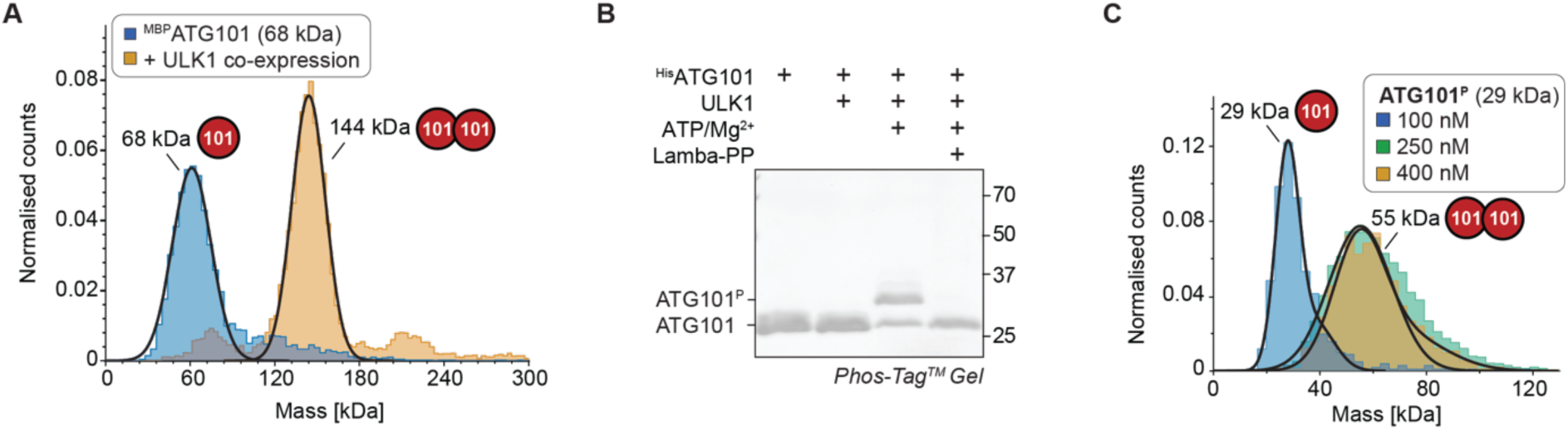
ULK1 activity induces ATG101 homodimerization. **a)** Co-expression of ULK1 kinase and ATG101 resulted in the stabilization of the ATG101 homo-dimer. Mass-photometry experiment showing size distribution of 250 nM purified ATG101 co-expressed with or without ULK1. **b)** ATG101 can be phosphorylated by purified ULK1. ATG101 was incubated with ULK1 (1:100 ratio) and Mg^2+^/ATP for 30 min at 27 °C. Phosphorylation was subsequently removed by the addition of lambda-phosphatase for 30 min at 27 °C. **c)** Phosphorylation stabilizes ATG101 homodimerization. Mass-photometry experiment showing size distribution at various concentrations of purified ATG101 *in vitro* phosphorylated by ULK1.

### Influence of ULK1 activity on ATG101 auto-activation dynamics

The results so-far have created a paradox: ATG101 homo-dimerization - despite being competitive with the ATG13-ATG101 complex, as well as weaker and transient – seems to boost heterodimer formation. We therefore wondered what the relevance and function of the ATG101 homo-dimerization could be. Given that the phosphorylation of ATG101 by ULK1 induces a structural conversion in ATG101, we wondered if it could influence the interaction kinetics of ATG101. When performing a pulldown experiment that monitors ATG13-ATG101 heterodimerization, we observed that phosphorylated ATG101 would interact stoichiometrically with ATG13 within a few minutes while untreated ATG101 did not (**Figure 4A,B** and **S4A**). Similar results were observed with the phospho-mimetic ATG101 mutants (**Figure 4C** and **S4B**). Next, we quantified the interaction kinetics of the ATG13-ATG101 complex using the FP assay. Using low concentrations of phosphorylated ATG101 (20 nM), the interaction with ATG13 was unchanged and slow (1.47 × 10^-5^ μM^-1^s^-1^) (**Figure S4C,D**). However, upon incrementally increasing the concentration of phosphorylated ATG101, we again observed a similar non-linear stark increase in ATG13-ATG101 complex assembly (**Figure 4D,E**). This observed effect was a similar acceleration as seen before using unphosphorylated ATG101 at elevated concentrations, but now occurred at much lower concentrations. Similarly, the ATG9A-ATG13-ATG101 complex forms within a few minutes when using the phosphorylated ATG101, while the untreated ATG101 does not allow prompt complex formation (**Figure 4F,G** and **S4E**). Similar results were obtained when using the phospho-mimetic ATG101 mutants (**Figure 4H** and **S4F**).

**Figure 4:**
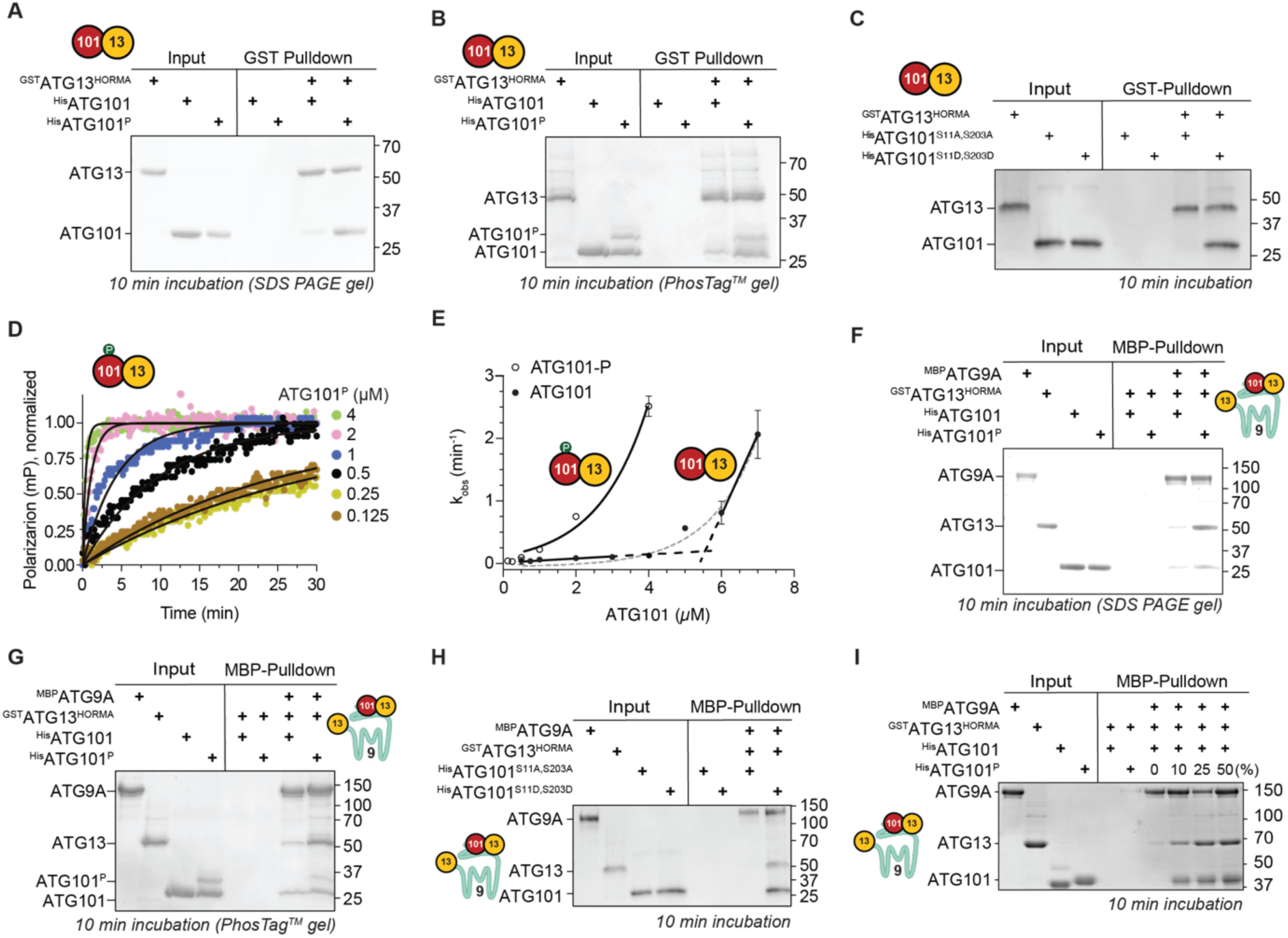
Activated ATG101 auto-catalytically promotes ATG9A-ATG13-ATG101 complex formation. **a,b)** Phosphorylation activates ATG101 to promote a quick interaction with ATG13. Panels **a** (SDS-PAGE) and **b** (Phos-tag^TM^ gel) show pull-down experiments after 10 min incubation of 0.2 μM GST-ATG101 as bait and 0.4 μM His-ATG101. Quantified in **S4A** from three independent experiments. **c)** Introducing phospho-mimetic mutants S11D and S203D in ATG101 facilitate prompt dimerization of ATG13-ATG101. Pull down experiment after 10 min incubation of 0.2 µM of ATG13 as bait and 0.4 µM ATG101 as prey. Quantification in **S4B** of three independent technical replicates. **d,e)** Phosphorylation activates ATG101 to promote a quick interaction with ATG13. Time zero is the first time point after mixing 50 μM unlabelled ATG13 with indicated phosphorylated ATG101^Alexa488^ concentrations. Shown in **d** are single measurements representative of at least three independent technical replicates of the experiment. After single exponential fitting of the curves, the apparent first order rate constants (k_obs_) were plotted as function of ATG13 concentration in **e**, with the results from panel **1e** shown as reference. **f,g)** Phosphorylation activates ATG101 to promote a quick assembly of the ATG9A-ATG13-ATG101 complex. Panels **f** (SDS-PAGE) and **g** (Phos-tag^TM^ gel) show pull-down experiments after 10 min incubation of 0.2 μM MBP-ATG9A as bait and 0.4 μM His-ATG101 and GST-ATG13, and the result is quantified in **S4E** from three independent experiments. **h)** Introducing phospho-mimetic mutants S11D and S203D in ATG101 facilitate quick assembly of the ATG9A-ATG13-ATG101 complex. Pull down experiment after 10 min incubation of 0.2 μM MBP-ATG9A as bait and 0.4 μM His-ATG101 and GST-ATG13 as prey. Quantification in **S4F** of three independent technical replicates. **i)** After being primed by phosphorylation, ATG101 will auto-catalytically activate other ATG101 molecules to promote quick association of ATG101 regardless of phosphorylation status. Pulldown experiment monitoring ATG13-ATG101 complex formation using varying relative amounts of phosphorylated ATG101 separated on a 15 % SDS-PAGE. ATG101 and ATG13 were mixed at a final concentration of 400 nM each. Quantification in **S4G** of three independent technical replicates.

Interestingly, despite the typical incomplete phosphorylation by ULK1, we invariably observed complete and rapid binding of both phosphorylated and unphosphorylated ATG101 to ATG13 and ATG9A (**Figure 4B,G**). Upon mixing 400 nM stoichiometric total amounts of ATG101 and ATG13, the presence of 10% of phosphorylated ATG101 sufficed to accelerate the assembly of the complex (**Figure 4I** and **S4G**). Overall, this shows that phosphorylation activates ATG101, and although phosphorylated ATG101 can interact with ATG13-ATG9, but is not a requirement for the interaction. After being primed by phosphorylation, ATG101 will auto-catalytically activate other ATG101 molecules to promote quick association of ATG101 regardless of phosphorylation status.

### Memory of activation of ATG101 is retained many hours after dephosphorylation

Presumably the activation of ATG101 is reversible in order to achieve a responsive system. Moreover, since phosphorylation is not strictly required for homo-dimerization nor the interaction with ATG13 or ATG9A, we wondered what would happen if we removed the phosphorylation. Therefore, we prepared four ATG101 samples, where we either left ATG101 untreated (A), purified ATG101 after a partial ULK1 treatment (B), or subsequently removed all phosphorylation using the generic lambda phosphatase and allowed ATG101 to revert to the inactivated state by waiting either 30 minutes (C) or overnight (D) (**Figure 5A**). We used these ATG101 samples to assess the kinetics of ATG9A-ATG13-ATG101 complex assembly by performing a pulldown experiment after 10 minutes after mixing them with separately purified ATG13 and ATG9A (**Figure 5B-D**). As before, no complex formed in this short time frame with the untreated ATG101 (A), while an incompletely phosphorylated ATG101 (B) would promote complete complex formation. Surprisingly however, although all phosphorylation was removed (samples C and D), ATG101 remained activated even after waiting overnight. Only the amount of ATG13 slowly reduced, yielding a stoichiometric 3:3:3 complex (**Figure 5D**).

**Figure 5:**
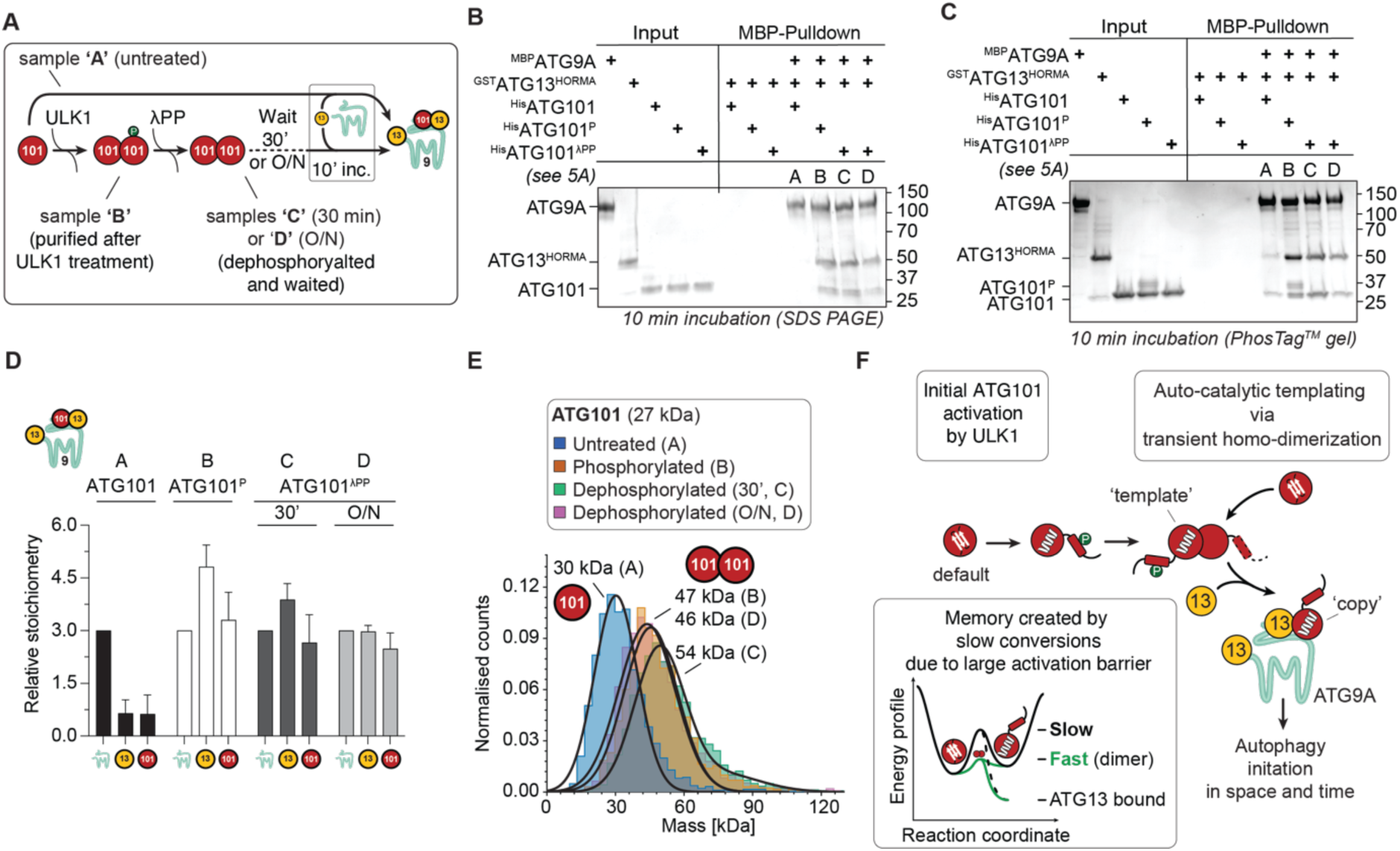
Memory of activation of ATG101 is retained many hours after dephosphorylation. **a)** Experimental design to test for the memory of activation after removal of phosphorylation of ATG101. **b-d)** Memory of activation of ATG101 is retained after dephosphorylation. Panels **b** (SDS-PAGE) and **c** (Phos-tag^TM^ gel) show pull-down experiments after 10 min incubation of 0.2 μM MBP-ATG9A as bait and 0.4 μM His-ATG101 and GST-ATG13. ATG101 treatment is described in panel **a**. Quantification from three independent experiments is shown in **d. e)** Dimerization of ATG101 is retained after dephosphorylation. Mass-photometry experiment showing size distribution of 200 nM purified ATG101 before and after phosphorylation by ULK1, and subsequent dephosphorylation. **f)** Model of the influence of ULK1 activity on ATG101 auto-activation dynamics, leading to quick assembly of the ATG9A-ATG13-ATG101 complex. Homologous to the MAD2-template model, ATG101 activation is likely auto-catalytically propagated by lowering activation energy of transition (see inset) allowing for ‘copying’ the state from the activated ‘template’ ATG101. This mechanism would support a responsive local initiation of autophagy.

Indeed, the homo-dimerization of ATG101 is persistent long after it had been dephosphorylated (**Figure 5E**). Overall, these experiments show that the phosphorylation is the trigger of the structural change that allows the dimerization and the increased interaction kinetics, is auto-catalytically propagated to non-phosphorylated ATG101 molecules. At this point, the phosphorylation becomes redundant for maintaining the structural change and homo-dimerization, yielding in a memory of activation as ATG101 ‘remembers’ its activation for many hours after dephosphorylation.

## Discussion

Following the induction of autophagy, a complex array of proteins promptly converges to create its initiation machinery. The ATG13-ATG101 dimer coordinates the recruitment of a variety of downstream factors to the autophagosome formation site^3,13,15,16,20–23^. Although some ATG13-ATG101 dimer can be found pre-assembled in fed cells, it only induces autophagy at limited levels^15^. It is likely that their interactions with its various binding partners, which might be mutually exclusive and have a multitude of functions, is regulated at multiple levels. This likely includes the regulation of the metamorphic behaviour of ATG13 and ATG101, as mutants in predicted metamorphic regions in ATG13 and ATG101 both do not allow for the assembly of the initiation machinery and show strong autophagic defects^13^. To shed light on the regulatory mechanisms that dictate complex assembly, it is pertinent that the rate-limiting steps are identified as these are ultimately what determine when and where the initiation machinery assembles. In this manuscript, we have shown that the enigmatic core autophagy protein ATG101 interacts exceedingly slowly with its interaction partners. ATG101 can spontaneously interact, but our quantifications show that it is 4-orders of magnitude slower than typical protein-protein interactions and therefore requires over 24 hours to complete at physiologically relevant concentrations. This timing is not conducive to responsive autophagosome formation, which is estimated to take 5 to 10 minutes in mammals^46,47^. We show that this unusual behavior of ATG101 is likely the result of the large activation energy required for the structural conversion of ATG101, where the most C-terminal beta-strand needs to be dislodged from an extended beta-sheet, and refolds as an alpha-helix. This structural metamorphosis, which is stabilized by transient homo-dimerization, is obligatory to convert from its default autoinhibited conformer to an active autophagy promoting conformer (**Figure 5F**). We postulate that this rate-limiting event provides an opportunity to regulate when and where autophagy is initiated. The kinase ULK1, a well-characterized interaction partner of ATG101 in the cell^2–5^, is the presumable candidate to perform this regulatory task, as we show that it phosphorylates ATG101 resulting in homo-dimerization of ATG101. Crucially, this dimerization induces the structural conversion of ATG101 which is essential to accelerate ATG101 activation dynamics. Further structural analysis will be required to confirm the (AlphaFold3) prediction that the position of the phosphorylation stabilizes ATG101 in the converted fold. Variations of this mechanism have been observed in related metamorphic HORMA domain proteins^48^, where kinases initiate the activation of otherwise inhibited signaling nodes by accelerating their structural conversion^27,28^. Once converted, these subsequently accelerate additional conversions via transient homo-dimerization^25,29–32^. This cascading ‘templating’ mechanism^26^ would thus be auto-catalytic and thereby efficiently inducing effector complex formation in a spatio-temporally controlled manner. The observation that only a minor portion of ATG101 needs to be phosphorylated to induce all molecules to interact with ATG9A and ATG13, strongly suggests that ATG101 also allows a similar auto-catalytic regulatory ‘templating’ mechanism (**Figure 5F**). The ATG101 homo-dimerization would lower the activation barrier by stabilizing a ‘reaction intermediate’ that catalyses the structural conversion and thus the interaction to ATG13 and ATG9A (**Figure 5F**, inset). We show that stabilization of homo-dimer is not directly dependent on phosphorylation, but rather activates the transition of ATG101 into a form that can homo-dimerize and interact with ATG13. The ATG101 homo-dimer cannot interact with ATG9A, securing the order of events: the transient nature of the homo-dimer promotes binding to ATG13 first and together they assemble the ATG9A-ATG13-ATG101 complex. This complex assembles the rest of the initiation machinery, where for example the newly ejected alpha-helix can serve as a receptor for further autophagy initiation factors, such as the PI3-kinase complex^22^. Regulating interactions via interfaces exclusively accessible after structural refolding would prevent premature recruitment of initiation factors, a mechanism that finds precedent in other HORMA domain proteins^49^, including ATG13^13^. Upon silencing autophagy, the initiation machinery is likely disassembled. The conversion of ATG101 back to the default inhibited state is equally slow, which constitutes the long memory of activation even after dephosphorylation (**Figure 5F**, inset). We expect that currently unknown external factors will accelerate this conversion to silence autophagy. Overall, this extra-ordinary mechanism would create responsive feedforward mechanism to control autophagosome biogenesis in time and space.

## Acknowledgements

We are grateful to Stefanie Asper for expert technical assistance. We thank the MPI of Multidisciplinary Sciences for support, and in particular the live-cell imaging facility (MPI-NAT) for access to the plate reader and Mass Photometer. We thank the Stein Lab (MPI-NAT) for sharing the SortaseA plasmid. We also acknowledge the help of Monica Raabe, Ralf Pflanz and Sabine König from the Proteomics Facility (MPINAT) for assistance with sample handling. We are grateful to the Griesinger laboratory (MPI-NAT) for access to the CD spectrometer. We thank the current members of the Faesen Lab for critical comments, discussions and helpful suggestions. Funding for this work was provided by the Max-Planck Society (to HG and HU) via the Max-Planck Research Group Leader program (to AF) and by the German Research Foundation (Deutsche Forschungsgemeinschaft; DFG) via SFB1190 (project P19 to AF) and FA 1752/3-1 (project number 542770124 to AF).

## Author contributions

AP, SS and ACF designed *in vitro* experiments and analyzed results. AP and SS set up recombinant expression systems, established purification protocols and purified proteins. AP and SS performed *in vitro* experiments. GN helped with the analysis of CD data and assisted with SESCA calculations and analysis. OD performed mass-spectrometry and analysis, supervised by HU. ACF supervised project and wrote the manuscript.

## Data and materials availability

All data are available in the main text or the supplementary materials.

## Materials and methods

### Expression and purification of proteins

**Table 1.** Buffers used for protein purification.

| Buffer | Composition |
| --- | --- |
| A | 150 mM NaCl, 20 mM HEPES pH 7.5, 5 %. Glycerol, 0.5 mM TCEP |
| B | 150 mM NaCl, 20 mM HEPES pH 7.5, 5 %. Glycerol, 0.5 mM TCEP, 0.03 % DDM |

#### ATG101 (insect cells)

ATG101 was cloned as an N-terminal GST, 6xHis-MBP, or Strep fusion construct from Hi5 cells using the biGBac expression system. 5 ml of V2 baculovirus was infected in 400-500 ml Hi5® cell cultures at 1×10E6/ml density. Cells were harvested at 72h of infection. Cell pellets were washed and resuspended in buffer A supplemented with 1 mM PMSF and 10 µg/50 mL DNAseI. The resuspended cells were lysed by sonication using the Branson Sonifier at an amplitude of 25 %, with a 5 min cycle at a pulse of 5 seconds on and 10 seconds off. The lysate was cleared by centrifugation at 15,000 rcf for 30 minutes. The cleared lysate was filtered using a 0.2 µM filter and passed over a 5mL GSTrap, MBPTrap, or StrepTrap column (Cytiva) for separation by affinity chromatography. Columns were pre-equilibrated with buffer A and unbound lysate after sample application with buffer A. Tagged ATG101 was then eluted in buffer A, supplemented with either 20 mM reduced glutathione at pH 7.0, 20 mM maltose, or 5 mM desthiobiotin depending on the column used. Eluted protein was concentrated in an Amicon-Ultra-15 Centrifugal Filter with a 10 kDa MWCO (Millipore). Sample was applied for further purification using Size Exclusion Chromatopgraphy on a Superdex 75 Increase 10/300 GL column equilibrated with buffer A. Peak fractions corresponding to ATG101 at the optimal retention volume were collected and again concentrated in an Amicon-Ultra-4 Centrifugal Filter 10 kDa MWCO. Protein aliquots were flash-frozen in liquid nitrogen and stored at -80 °C.

#### ATG101 (E. coli)

ATG101 with an N-terminal 6xHis Tag fusion was produced in E. coli cells. 100 µg plasmid was transformed in 50 µL Rosetta2 cells using heat-shock. Cells were grown in 500 ml cultures to reach an O.D. of 0.6 after which 0.2 mM IPTG was added. Cells were grown further for 16 h at 16 °C and harvested by centrifugation. Downstream purification steps remain the same as above except using a 5 mL HiTrap column for affinity chromatography. Column equilibration and unbound lysate washing was carried out using buffer A supplemented with 10 mM Imidazole. Tagged protein was eluted using buffer A supplemented with 250 mM Imidazole. Size Exclusion Chromatography and storage of protein was as above.

#### ATG9A

ATG9A with an N-terminal 6xHis-MBP tag was expressed in Hi5 insect cells using the biGBac expression system^50,51^. Cell pellet was resuspended in lysis buffer A supplemented with 1% DDM, 1 mM PMSF and 10 µg/50 mL DNAseI. Cells were lysed on ice for at least 1 h by stir-mixing. The lysate was then diluted 3 times with DDM-free buffer A and stir-mixed for another 30 min before clarifying by centrifugation at 15,000 rcf for 30 min. Columns were pre-equilibrated with buffer A and unbound lysate after sample application with buffer A. Tagged ATG9 was eluted in buffer A, supplemented with 20 mM maltose. Peak fractions were concentrated using a n Amicon-Ultra-15 Centrifugal Filter with a 100 kDa MWCO (Millipore) followed by size exclusion chromatography using a Superose 6 10/300 or 16/600 column (Cytiva) pre-equilibrated with buffer B. The purified protein was concentrated using an Amicon-Ultra-4 Centrifugal Filter with a 100 kDa MWCO (Millipore), flash-frozen in liquid nitrogen, and stored at - 80 °C.

#### ATG13

ATG13 full-length and HORMA (1-200) constructs were cloned as N-terminal GST or 6xHis-MBP fusion constructs from Hi5 cells using the biGBac expression system. 5 ml of V2 baculovirus was infected in 400-500 ml Hi5® cell cultures at 1×10E6/ml density. Cells were harvested at 72 h of infection. Cell pellets were washed and resuspended in buffer A supplemented with 1 mM PMSF and 10 µg/50 mL DNAseI. The resuspended cells were lysed by sonication using the Branson Sonifier at an amplitude of 25 %, with a 5 min cycle at a pulse of 5 s on and 10 s off. The lysate was cleared by centrifugation at 15,000 rcf for 30 minutes. The cleared lysate was filtered using a 0.2 µM filter and passed over a 5mL GSTrap, MBPTrap, or StrepTrap column (Cytiva) for separation by affinity chromatography. Columns were pre-equilibrated with buffer A and unbound lysate after sample application with buffer A. Tagged ATG13 was then eluted in buffer A, supplemented with either 20 mM reduced glutathione at pH 7.0 or 20 mM maltose depending on the column used. Eluted protein was concentrated in an Amicon-Ultra-15 Centrifugal Filter with a 10 kDa MWCO (Millipore). Sample was applied for further purification using Size Exclusion Chromatography on a Superose 6 Increase 10/300 GL column equilibrated with buffer A. Peak fractions corresponding to ATG13 at the optimal retention volume were collected and again concentrated in an Amicon-Ultra-4 Centrifugal Filter 10 kDa MWCO. Protein aliquots were flash-frozen in liquid nitrogen and stored at - 80 °C.

#### ULK1

ULK1 with an N-terminal GST-tag fusion was expressed in Hi5 cells using the biGBac expression system. Harvested cells were lysed by stir-mixing in buffer A supplemented with 1% DDM, 2 mM PMSF, 100 µM Leupeptin for 30 min - 1 h on ice. The lysate was then diluted 3 times with buffer A and clarified by centrifugation at 15,000 rcf for 30 min at 4 °C. The protein was purified from the lysate by affinity chromatography using a 5 mL GSTrap column (Cytiva) pre-equilibrated with buffer A and eluted using buffer A supplemented with 20 mM reduced glutathione at pH 7.0, as elution buffer. The purified protein was concentrated, flash-frozen in liquid nitrogen and stored at -80 °C.

#### GST, MBP and Strep pulldown assays

GST, MBP and Strep pulldown experiments were performed using GSH Sepharose beads, MBP beads (Amylose resin) (NEB), or StrepTactin Superlow Plus (Qiagen) beads pre-equilibrated by washing in buffer B. Concentrations of proteins mentioned in the figure legends were incubated separately before being added to 20 µL of corresponding bead resin. Beads were spun down at 500 rcf for 15 s. The supernatant was removed, and beads were washed twice with 300 μL buffer, with each wash & spin-down cycle not more than 30 s. The supernatant was removed completely, and 10µL of 4x Laemmli sample loading buffer was added to the beads. Samples were run on either 12 % SDS–PAGE gel made in house or a 4-15 % TGX-gel (BioRad). Bands were visualized with Coomassie Brilliant Blue (CBB) staining. Detailed information regarding concentration, temperature, and other variations in the pulldown is provided in the figure legends.

#### Cross-linking Mass Spectrometry

To cross-link the ATG101 dimer, two approaches were taken. First, 5 µM of ATG101 was pre-incubated for 1 h at 20 °C, followed by a 30-min incubation at 20 °C with the indicated amount of BS3 (Thermo Scientific), and subsequent quenching by addition of 1M Tris-HCl, pH 7.0 (final concentration 25 mM) for 15 min. Second, 2 µM of MBP-ATG101 was incubated for 1 h at 20 °C with 6 µM of Strep-ATG101. The formed dimer-complex was pulled down and washed to remove excess unbound protein (as in the procedure for *in vitro* pull-down assay described above). Proteins were then separated by SDS-PAGE using a 4–12 % gradient gel (BioRad).

Upshifted bands corresponding to the cross-linked complexes were excised from the gel and in-gel digested with trypsin (Shevchenko et al.et al., 2006). In brief, samples were reduced with 10 mM dithiothreitol and alkylated with 55 mM iodoacetamide and subsequently digested with trypsin (sequencing grade, Promega) at 37 °C for 18 h. Extracted peptides were dried in a SpeedVac vacuum concentrators (Thermo Scientific) and dissolved in a buffer composed of 5 % acetonitrile and 0.1 % trifluoroacetic acid. Samples were analyzed by liquid chromatography electrospray ionization mass spectrometry (LC-ESI-MS). For this, peptides were online separated on a Dionex UltiMate 3000 UHPLC system (Thermo Scientific) equipped with an in-house-packed analytical C18 column (75 µm × 300 mm, ReproSil-Pur 120 C18-AQ, 3 µm, Dr. Maisch GmbH) and coupled to an Exploris 480 mass spectrometer (Thermo Scientific). The latter was operated in data-dependent mode with the following settings: MS1 resolution, 120 000, normalized AGC target, 300%, scan range 350-1550 m/z; cycle time 3 s; MS2 resolution, 30 000, normalized collision energy for HCD fragmentation, 30 %, maximal injection time, 128 ms, normalized AGC target, 75%, isolation window, 1.6. Only precursors with charge state of 3-8 were selected for MS2. To verify protein composition of the samples, Thermo raw files were searched first with MaxQuant 2.6.7.0 against a database encompassing UniProt reference proteomes of Homo sapiens and Trichoplusia ni as well as the sequences of MBP- and Strep-tagged ATG101 and protein contaminants commonly found in MS samples. Cross-linked peptides were identified in a subsequent search using pLink3.0.16 46 and a database containing only trypsin, MBP- and Strep-tagged ATG101 sequences. FDR was set to 1% at spectral level. The crosslinks were visualized using xiNET47.

#### Sortase-mediated fluorescence labelling of ATG101

To make a fluorescently labelled peptide for enzymatic conjugation by SortaseA, 1 mol GGGC tetrapeptide (dissolved in 100 % DMSO, ordered from GenScript) was mixed with 2 mol Alexa-488® C-5 Maleimide dye (dissolved in 100 % DMSO, ThermoFisher^TM^ scientific) for 2 h at 20 °C in a dark environment. The reaction was quenched by adding 10 mM DTT. The resulting labelled peptide GGGC^-Alexa488^ was flash-frozen stored at -80 °C.

To fluorescently label ATG101, 30-40 µM of purified ATG101 in buffer A was mixed with 1 µM SortaseA enzyme (purified in house), 10 mM CaCl_2_ and 150 µM GGGC-^Alexa488^ labelled peptide. The reaction mixture was incubated at 27 °C for 1-2 h in a thermoshaker. Labelled protein obtained was separated from enzyme and excess peptide by SEC with UV-490nm monitoring. Absorbance of collected protein was measured at 280 and 490 nm using NanoDrop^TM^ one to determine labelling efficiency. Labelled protein was flash-frozen and stored at -80 °C for further use.

#### In vitro phosphorylation of ATG101

For phosphorylation by ULK1, ATG101 purified from E. coli was used as substrate. In-house purified ULK1 at a final concentration of 3-5 µM, was mixed with ATG101 in buffer A supplemented with 0.5 mM ATP and 10 mM MgCl_2_ and incubated at 25 °C for 30 min. Phosphorylated ATG101 was separated from ULK1 and excess ATP by SEC. Phosphorylated protein was flash-frozen and stored at -80 °C for further use. Analysis of phosphorylated protein by gel electrophoresis was either done by separation on SuperSep^TM^ PhosTag (Fujifilm) gels or normal 12 or 15 % SDS-PAGE gels as mentioned in the captions, followed by staining with Pro-Q Diamond^TM^ Phosphostain (Thermo Fisher) as per manufacturer’s protocol.

#### Mass Spectrometry to confirm phosphorylation sites

ATG101 purified from E.coli was phosphorylated *in vitro* using ULK1 kinase as described below. The samples further purified by SEC for kinase and ATP removal were loaded on a 12% SuperSepTM PhosTag12.5% gel (Fig.). The upshifted band signifying phospho-ATG101 was checked to confirm previously known ULK1 phosphorylation sites by mass spectrometry. For this, in-gel digestion was performed as described above for cross-linked samples followed by an enrichment of phospho-peptides using Titanium Dioxide TopTips (Glygen) and a subsequent LC-ESI-MS analysis. MASCOT 2.3.02 was used as a search engine. Carbamidomethylation of cysteine was set as fixed modification; oxidation of methionine and phosphorylation of serine/threonine/tryptophan was considered as variable modifications. Peptides corresponding to Ser11 and Ser203 are shown in Supplementary Table **S2**.

#### Fluorescence polarization assay

All FP-based binding measurements were performed using the BioTek Synergy Neo2 microplate reader (Agilent). For kinetic experiments measuring association rates, labelled ATG101 (or phosphorylated ATG101) or labelled ATG13-101 was prepared in buffer B in concentrations indicated in the figure legends and added to a 96-well assay plate. ^MBP^ATG9A or ^MBP^ATG13 full-length at concentrations indicated in the figures was added into ATG101 or ATG13-101 containing wells and measurements recorded. The final reaction volume in each well was 100 µL. For longer measurements, a time course was set for 12 h, readings taken every 45 s. Gain was set to 50/50. For shorter measurements, a time course was set for 30 min or 1 h, readings taken every 10 s. Gain was set to 50/50. Fluorescence polarization was selected as the readout, using filter cubes Dual FP (top) 480/520 (bottom) (Agilent). Temperature at the time of all recordings was 22 °C ± 1 °C. Data analysis for kinetic experiments was performed by fitting obtained curves to a one-phase association equation using GraphPad Prism ®. Observed rate contstants *k*_obs_ from this equation were plotted against the concentrations to fit into a linear regression. Association rates k_on_ were given by the slope of the equation in units µM^-1^ h^-1^.

For end-point saturation measurements, 10 nM of labelled ATG101 or 5 nM of ATG13-101 was pre-incubated for 16 h with serial dilutions of ^MBP^ATG9A or ^MBP^ATG13 full-length. An end-point measurement of fluorescence-polarization was performed using the same gain settings and filter cubes as mentioned above. Data obtained were plotted against the concentrations and fitted with a one-phase specific binding equation using GraphPad Prism ® to obtain the equilibrium dissociation constants (K_d_).

#### SDS-PAGE densitometric quantification

For SDS-PAGE quantification of relative stoichiometries, the density of each protein band scanned by a gelscanner (Epson) was calculated using GelAnalzyer (version 19). Densities were divided by the molecular weight of the protein and ratios were calculated by normalizing to the bait protein. Change in stoichiometries were visualized as bar plots. The results (n=3) are shown as mean + standard deviation calculated with GraphPad Prism (version 9.0.0).

#### Mass Photometry

Mass Photometry measurements were performed using a OneMP mass photometer (Refeyn Ltd, Oxford, UK). Data was acquired using the AcquireMP software (Refeyn Ltd. v2.3). For measurement, a drop of immersion oil was applied to the objective lens. Silicon gasket wells to hold the samples were then fixed on a clean dust-free cover slip placed on the lens stage. To find focus, filtered and degassed buffer A was pipetted into one gasket well and the focal point locked. Each sample at concentrations indicated in figure legends were pipetted add mixed nto a gasket well, and data were acquired with an acquisition time varying between 30 – 60s. The timing was adjusted to get a good number of landing events while avoiding saturation. DiscoverMP software (Refeyn Ltd. v2.3) was used to analyze the data.

#### CD spectrometry and prediction of secondary structures

All CD measurements were performed using the JASCO J815 spectrometer (JASCO inc.). 4 µM protein in buffer A was added to a rectangular quartz cell cuvette (0.1 cm pathlength) (JASCO inc.). A buffer blank was recorded before each measurement and automatically subtracted from the sample measurement. Spectra were recorded in a wavelength range of 260 – 190 nm with a data pitch of 1 nm. The scanning mode was set to continuous with a speed of 50 nm/min. Three repeats were collected for each spectrum. SESCA v097 was used for the analysis of the spectra. Before estimating the percentage of secondary structures, the spectrum was processed using SESCA process. Only data points within wavelengths of 250nm-205nm were chosen due to voltage limitations and the CD spectrum was converted to 1000 mean residue ellipticity units. SESCA Bayes was used for the structure estimations using the DS-dT basis set (1500 iterations where performed and a fraction of 0.1 was discarded). Heatmaps were generated using SESCA projection from the Bayesian estimations.

#### Bacterial and virus strains

|  |  |  |
| --- | --- | --- |
| DH10EMBacY™ | Geneva Biotech | Cat#10361012 |
| NEB®5-alpha | New England Biolabs | Cat#C2987H |
| Rosetta2™ | Novagen | Cat#71402 |

#### Cell lines

|  |  |  |
| --- | --- | --- |
| Sf9 insect cells<br>(*Spodoptera frugiperda*) | Invitrogen | Cat#10503433 |
| High Five insect cells<br>(*Trichopulsia Ni*) | Invitrogen | Cat#10747474 |

#### Culture media

|  |  |  |
| --- | --- | --- |
| LB medium | BioChemica | Cat#23143289 |
| LB-agar | Roth | Cat#X969.2 |
| ESF 921 medium | Expression Systems | Cat#96-001-01 |
| Sf-900 medium | Gibco | Cat#12658-019 |

#### Chemicals, peptides, and recombinant proteins

|  |  |  |
| --- | --- | --- |
| Dodecyl-β-D-maltoside (DDM) | Carl Roth | Cat#69227-93-6 |
| Triton X-100 | Merck | Cat#9036-19-5 |
| Amylose resin | NEB | Cat#E8021L |
| Pierce™ Glutathione Agarose | Thermo Fisher Scientific | Cat#16101 |
| Strep-Tactin Superflow Plus | IBA LifeSciences | Cat#30004 |
| SERVA FastLoad 1 kb DNA Ladder | SERVA | Cat#39317.01 |
| SERVA DNA Stain G | SERVA | Cat#39803 |
| Protein Dual Color Standards | BioRad | Cat#1610374 |
| Gel Loading Dye, Purple (6X) | NEB | Cat#B7024S |
| BS<sup>3</sup> (Bis(sulfosuccinimidyl)suberat) | Thermo Fisher Scientific | Cat#21580 |
| Maltose Monohydrat | Merck | Cat#6363-53-7 |
| L-Glutathion | Merck | Cat#70-18-8 |
| X-TREMEGENE 9 DNA Tranfection | Merck | Cat#6365779001 |
| 4–15% Mini-PROTEAN® TGX™ Precast Protein Gels | BioRad | Cat#4561083 |
| 96-well microplate (non-binding) | Corning CoStar | Cat#655906 |
| 0.45um Syringe Filter | AMSTAT | Cat#C0000629 |
| Amicon® Ultra-15 concentrator 100 kDa | Milipore | Cat#UFC910024 |
| Amicon® Ultra-15 concentrator 10 kDa | Milipore | Cat#UFC903024 |
| T5 exonuclease | Epicentre | Cat#T5E4111K |
| BSA | Roth | Cat#8076 |
| Phusion® High-Fidelity DNA Polymerase | NEB | Cat#M0530 |
| Q5® High-Fidelity DNA Polymerase | NEB | Cat#M0491 |
| Taq DNA Polymerase | NEB | Cat#M0267 |
| Taq DNA Ligase | NEB | Cat#M0208 |
| BamHI | NEB | Cat#R0136S |
| HindIII | NEB | Cat#R0104S |
| Benzonase® | Merck | Cat#E1014 |
| DpnI | NEB | Cat#R0176S |
| Alexa Fluor™ 488 C5 Maleimide dye | Invitrogen | Cat#A10235 |
| GGGC tetrapeptide | GenScript | Custom order |
| Calcium Chloride | Roth | Cat#275P.1 |
| IPTG | Roth | Cat#2316.5 |
| SuperSep™ Phos-Tag Precast gel | FujiFilm | Cat#193-16571 |
| Pro Q™ Diamond Phosphoprotein Gel Stain | Thermo Fisher Scientific | Cat#P33300 |

#### Recombinant DNA

|  |  |
| --- | --- |
| pColi-6x-His-ATG101-LPETGG | This study |
| pLIB-2xStrepII-ATG101-LPETGG | This study |
| pColi-6x-His-ATG101-L29R-H30R-LPETGG | This study |
| pLIB-6x-His-MBP-ATG9 | Nguyen, Lugarini et al., 2023 |
| pBIG-6x-His-MBP-ATG13<sup>HORMA</sup>-ATG101 | Nguyen, Lugarini et al., 2023 |
| pBIG-GST-ATG13<sup>HORMA</sup>(1-200) -ATG101 | Nguyen, Lugarini et al., 2023 |
| pLIB-6x-His-MBP-ATG13 | Nguyen, Lugarini et al., 2023 |
| pLIB-GST-ULK1 | Nguyen, Lugarini et al., 2023 |
| pLIB-6x-His-MBP-ATG13<sup>HORMA</sup> (1-200) | Nguyen, Lugarini et al., 2023 |
| pLIB-GST- ATG13<sup>HORMA</sup> (1-200) | Nguyen, Lugarini et al., 2023 |
| pLIB-6x-His-MBP-ATG101 | Nguyen, Lugarini et al., 2023 |
| pLIB-GST-ATG101 | Nguyen, Lugarini et al., 2023 |

#### Commercial assay kits

|  |  |  |
| --- | --- | --- |
| QIAprep Spin Miniprep Kit | Qiagen | Cat#27106 |
| Plasmid Midiprep Kit | Invitrogen | Cat#K210015 |
| QIAquick PCR purification Kit | Qiagen | Cat#28106 |

#### Software and algorithms

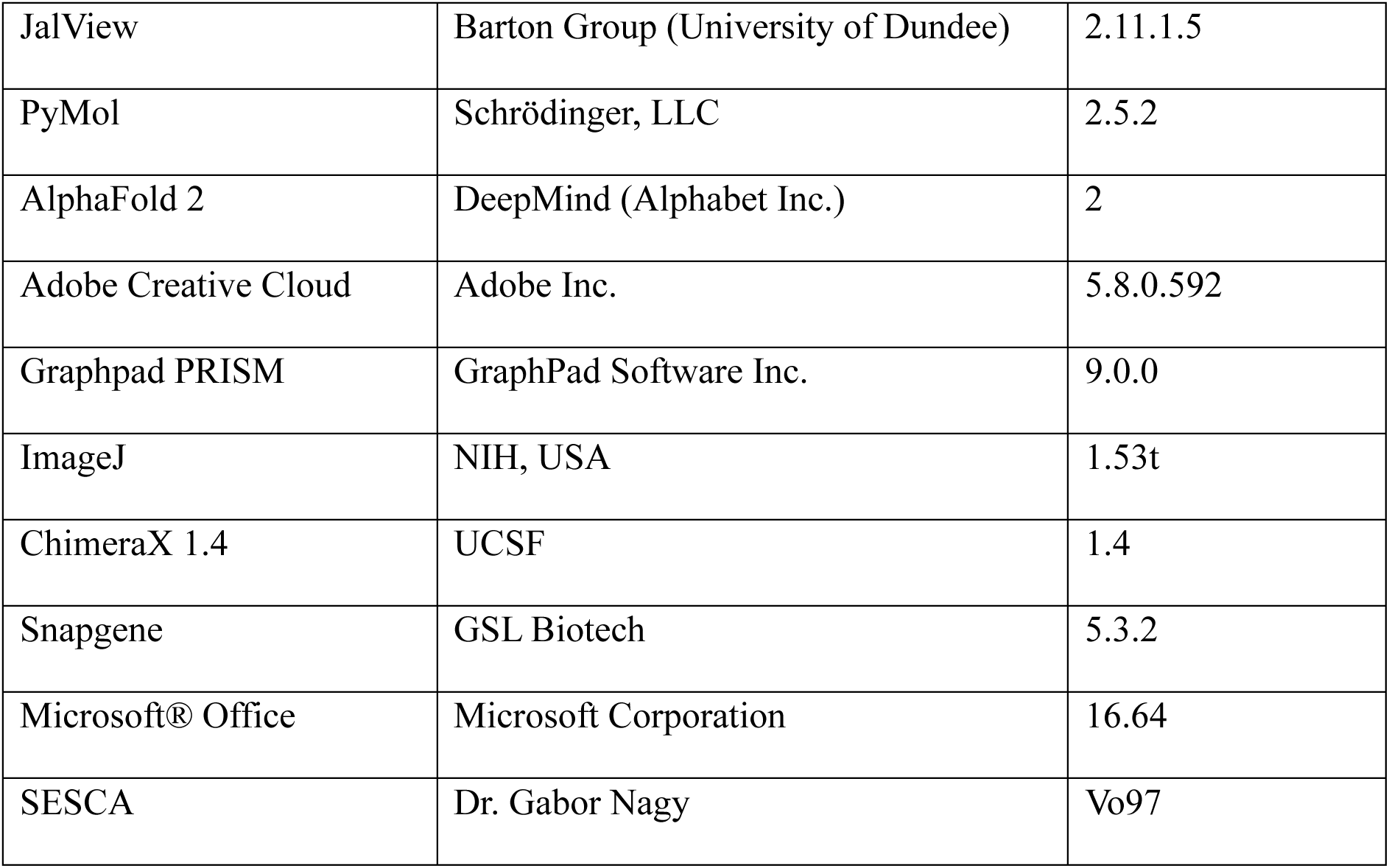
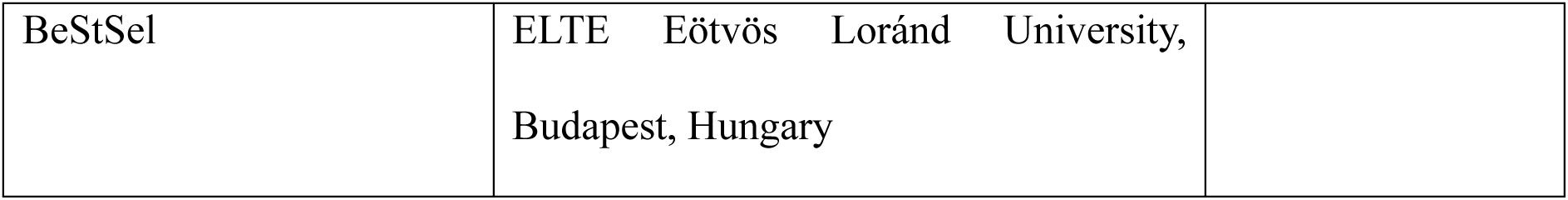

|  |  |  |
| --- | --- | --- |
| JalView | Barton Group (University of Dundee) | 2.11.1.5 |
| PyMol | Schrödinger, LLC | 2.5.2 |
| AlphaFold 2 | DeepMind (Alphabet Inc.) | 2 |
| Adobe Creative Cloud | Adobe Inc. | 5.8.0.592 |
| Graphpad PRISM | GraphPad Software Inc. | 9.0.0 |
| ImageJ | NIH, USA | 1.53t |
| ChimeraX 1.4 | UCSF | 1.4 |
| Snapgene | GSL Biotech | 5.3.2 |
| Microsoft® Office | Microsoft Corporation | 16.64 |
| SESCA | Dr. Gabor Nagy | Vo97 |

|  |  |
| --- | --- |
| BeStSel | ELTE Eötvös Loránd University,<br>Budapest, Hungary |

#### Columns and resins used for chromatography

|  |  |  |
| --- | --- | --- |
| MBPTrap™ HP (1 ml; 5 ml) | Cytiva | Cat#29048641;<br>Cat#28918778 |
| HiTrap TALON crude (1 ml; 5ml) | Cytiva | Cat#29048565;<br>Cat#28953766 |
| GSTrap™ HP (1 ml; 5 ml) | Cytiva | Cat#17528101;<br>Cat#17528201 |
| StrepTrap HP (5ml) | Cytiva | Cat#28907546 |
| HiLoad 16/600 Superdex 75 pg | Cytiva | Cat#28989333 |
| HiLoad 16/600 Superdex 200 pg | Cytiva | Cat#28989335 |
| HiLoad 16/600 Superose 6 pg | Cytiva | Cat#29323952 |
| Superdex 75 Increase 10/300 GL | Cytiva | Cat#29148721 |
| Superdex 200 Increase 5/150 GL | Cytiva | Cat#28990945 |
| Superose 6 Increase 10/300 GL | Cytiva | Cat#29091596 |
| Superose 6 Increase 5/150 GL | Cytiva | Cat#29091597 |

## Supplemental figures

**Figure S1:**
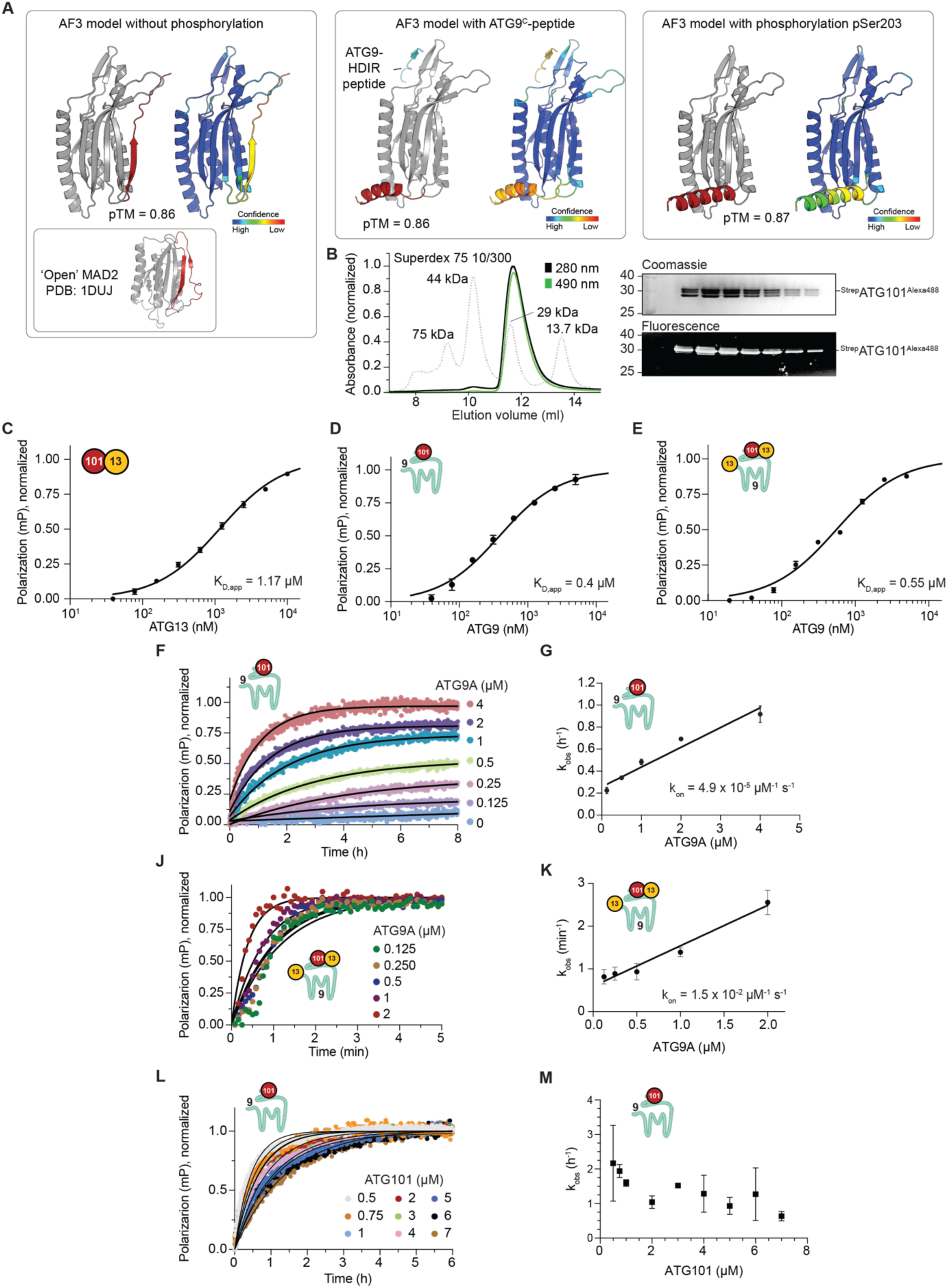
The interaction of ATG101 with ATG13 and ATG9A is exceedingly slow (related to figure 1). **a)** AlphaFold3 predicted models of ATG101 (left panel), with the ATG9-HDIR peptide (centre) and with Ser203 phosphorylation (right). Confidence of the predictions are indicated by respective pTM scores. Subset on the bottom left of the first panel shows the ‘open’ conformation of MAD2. **b)** Size exclusion chromatogram and SDS-PAGE of fluorescently labelled ATG101 used in the FP sensor throughout this study. **(c-e)** Fluorescence anisotropy-based saturation binding experiments of interactions within the ATG9A-ATG13-ATG101 complex. Cartoons on the top left of each panel indicate the interaction measured. 10 nM of ATG101^Alexa488^ (in **c** and **d**) or ATG13-ATG101Alexa^488^ (**e**) were incubated with varying concentrations of ATG13 or ATG9A. Equilibrium dissociation constants (K_d_) for each reaction were obtained by fitting a one-site specific binding equation in Graphpad Prism. **f,g)** The interaction of ATG101 with ATG9 is slow. Time zero is the first time point after mixing 20 nM ATG101^Alexa488^ with indicated ATG9A concentrations. Experiment is a single measurements representative of at least three independent technical replicates of the experiment. After single exponential fitting of the curves in **f**, the apparent first order rate constants (k_obs_) were plotted as function of ATG9A concentration in **g**, with k_on_ being the slope of the resulting line. **j,k)** Preformed ATG13-ATG101 complex interacts 3-orders of magnitude faster with ATG9A. Time zero is the first time point after mixing 10 nM ATG101^Alexa488^–ATG13 with indicated ATG9A concentrations. Experiment is a single measurements representative of at least three independent technical replicates of the experiment. After single exponential fitting of the curves in **j**, the apparent first order rate constants (k_obs_) were plotted as function of ATG9A concentration in **k**, with k_on_ being the slope of the resulting line. **l,m)** The interaction of ATG101 to ATG9A is progressively inhibited at higher ATG101 concentrations. Time zero is the first time point after mixing 50 µM ATG9A with indicated ATG101 concentrations. Experiment is a single measurements representative of at least three independent technical replicates of the experiment. After single exponential fitting of the curves in **l**, the apparent first order rate constants (k_obs_) were plotted as function of ATG9A concentration in **m**.

**Figure S2:**
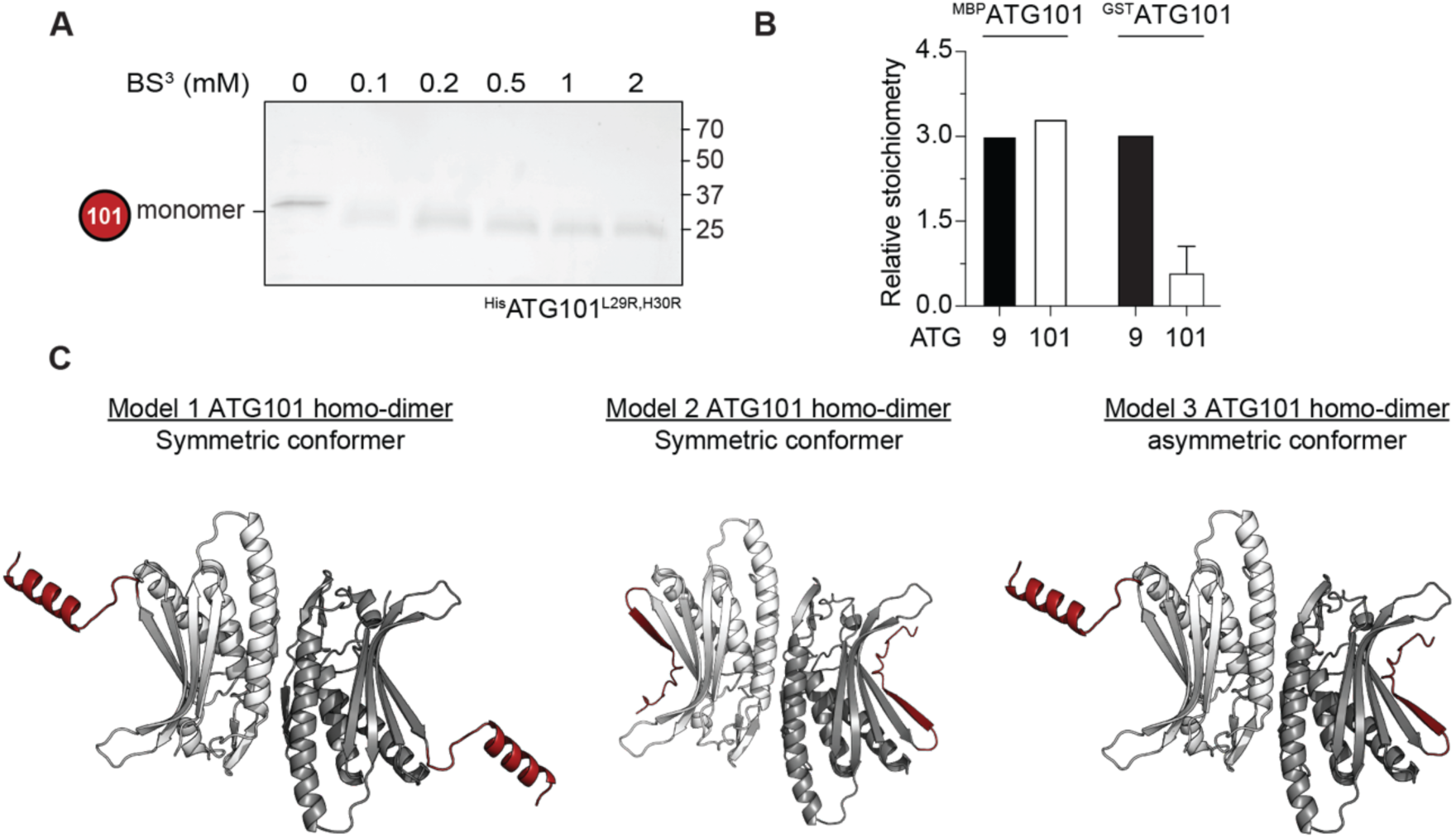
ATG101 transiently homo-dimerizes (related to figure 2). **a)** Incubation of ATG101 mutated in the dimer interface (L29R H30R; reference ^41^) with cross-linker BS3 does not create in proteins larger then monomeric ATG101. Input is shown in lanes with no BS3 (0 mM). **b)** Quantification of three independent technical replicates of the experiment in **2C**. **c)** Proposed models of conformationally symmetric or asymmetric ATG101 homo-dimers generated using AlphaFold3 and HADDOCK 2.4.

**Figure S3:**
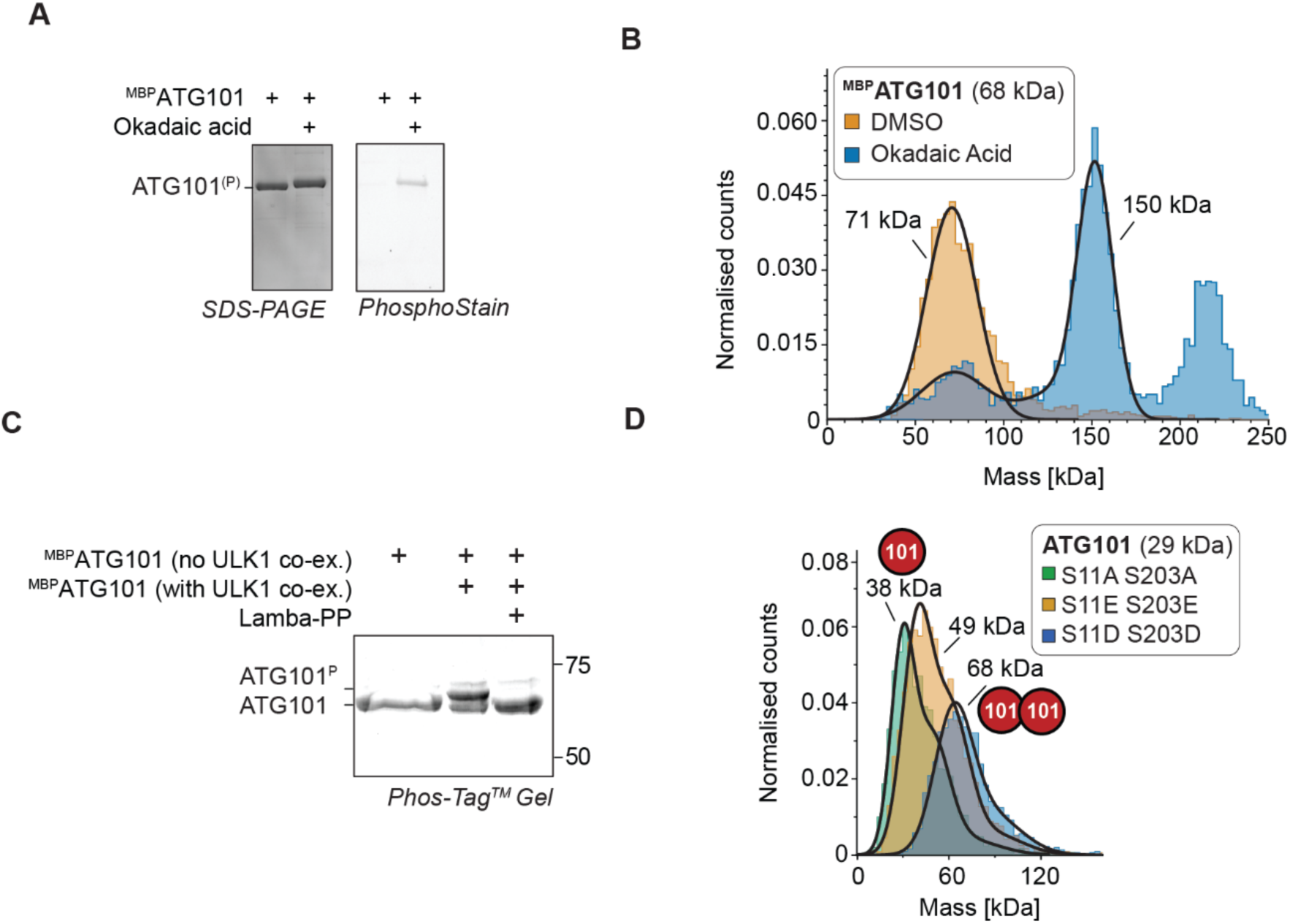
Phosphorylation by ULK1 initiates ATG101 homo-dimerization (related to figure 3). **a)** Purified ATG101 is phosphorylated if okadaic acid is added during the last few hours of the insect cell expression culture. **b)** Mass Photometry experiment showing the dimerization of purified ATG101 (150 nM) upon adding Okadaic acid to the insect cell expression culture. **c)** Phos-tag^TM^ gel showing that ATG101 can be reversibly phosphorylated by ULK1 co-expression. **d)** Mass Photometry experiment showing the effect of mutating the ULK1 phosphorylation sites in ATG101 (150 nM).

**Figure S4:**
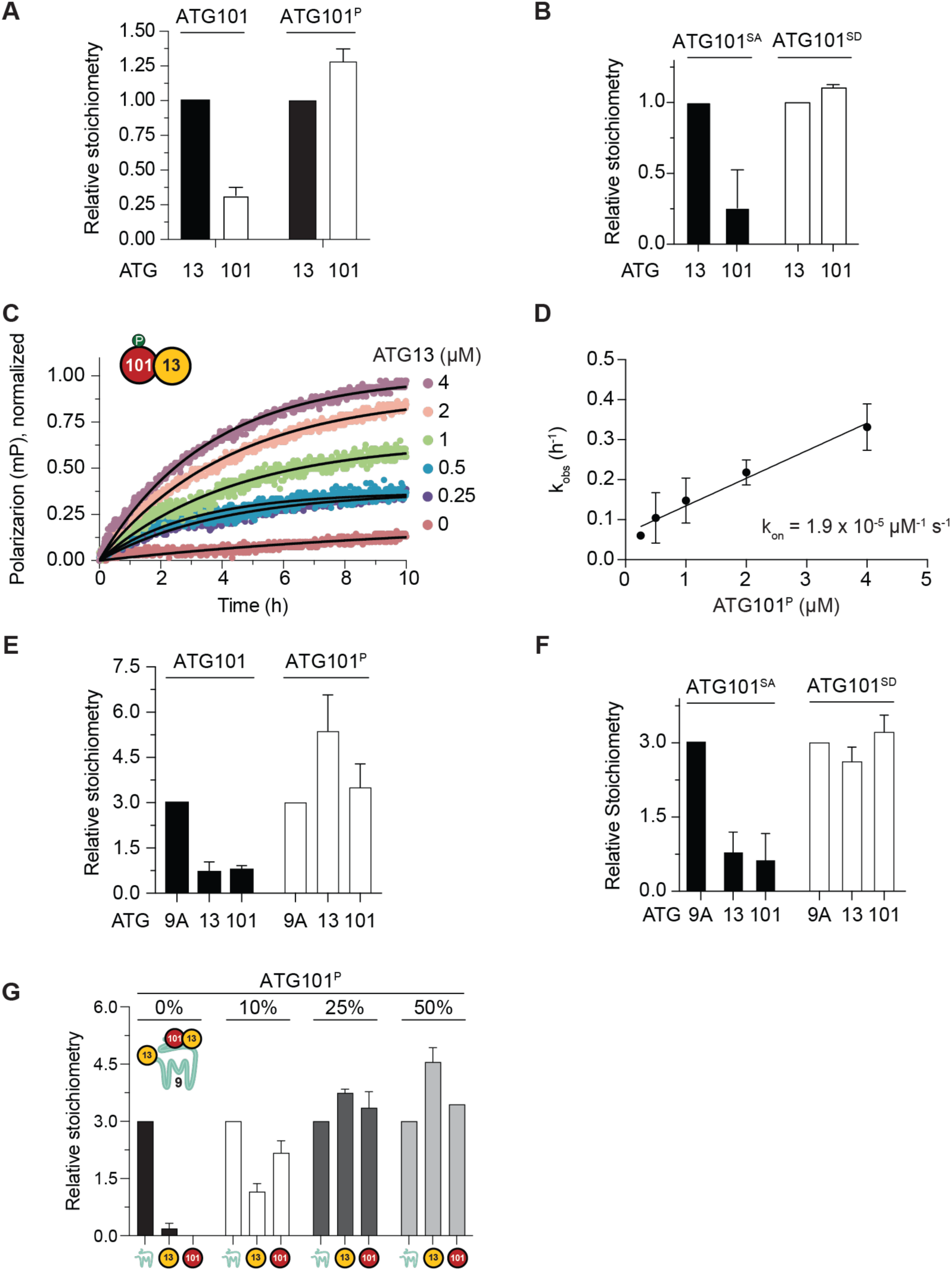
Activated ATG101 auto-catalytically elicits ATG9A-ATG13-ATG101 complex formation (related to figure 4). **a,b)** Quantification of three independent technical replicates of the experiment in **4A** and **4C**, respectively. **c,d)** Interaction of phosphorylated ATG101 at low concentrations with ATG13 is still exceedingly slow. Time zero is the first time point after mixing 20 nM phosphorylated ATG101^Alexa488^ with indicated ATG13 concentrations. Time-dependent changes in Fluorescence Anisotropy signal are single measurements representative of at least three independent technical replicates of the experiment. After single exponential fitting of the curves in **c**, the apparent first order rate constants (k_obs_) were plotted as function of ATG13 concentration in **d**, with k_on_ being the slope of the resulting line. **e-g)** Quantification of three independent technical replicates of the experiment in **4F**, **4H**, and **4I**, respectively.

**Supplementary Table 1.** List of crosslinks found in ATG101.

|  | Crosslinked Residue 1 | Crosslinked Residue 2 |
| --- | --- | --- |
| 1 | 36 | 78 |
| 2 | 36 | 143 |
| 3 | 40 | 143 |
| 4 | 78 | 36 |
| 5 | 84 | 107 |
| 6 | 107 | 84 |
| 7 | 107 | 147 |
| 8 | 143 | 36 |
| 9 | 143 | 40 |
| 10 | 143 | 143 |
| 11 | 143 | 147 |
| 12 | 147 | 107 |
| 13 | 147 | 143 |
| 14 | 147 | 147 |

**Supplemental Table 2:**
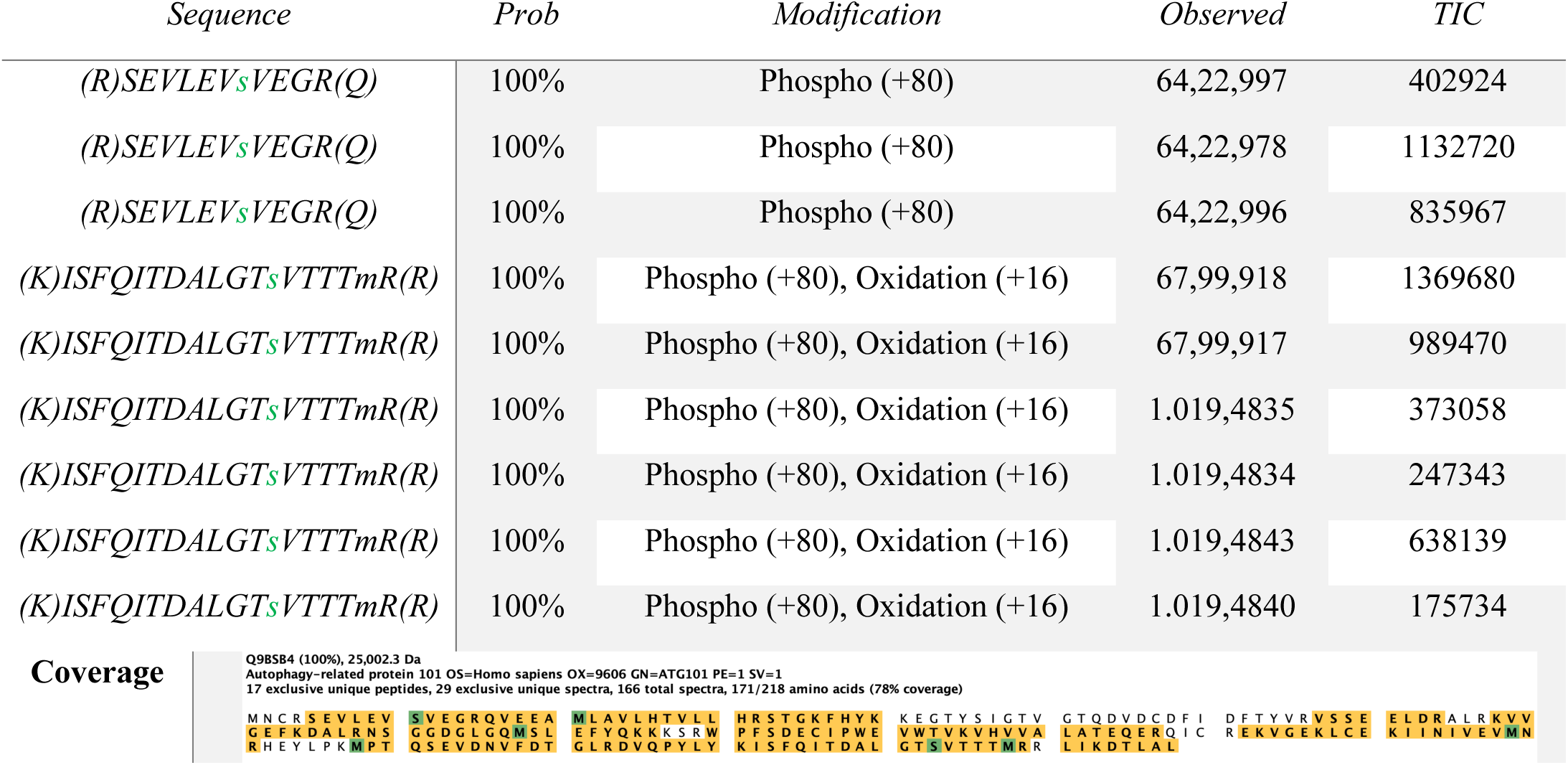
Phosphorylation sites of ATG101 by ULK1.

